# Modulation and recruitment of TRF2 at viral telomeres during human herpesvirus 6A/B infection

**DOI:** 10.1101/514075

**Authors:** Shella Gilbert-Girard, Annie Gravel, Vanessa Collin, Darren J. Wight, Benedikt B. Kaufer, Eros Lazzerini-Denchi, Louis Flamand

## Abstract

Human herpesviruses 6A and 6B (HHV-6A/B) can integrate their genomes into the telomeres of host chromosomes. The HHV-6A/B genomes contain telomeric repeats essential for integration. Whether HHV-6A/B infections impact telomere homeostasis remains to be studied. We report that during infection, a massive increase in telomeric signals is observed. Such telomeric signals are detected in viral replication compartments (VRC) that colocalize with the viral IE2 and P41 proteins. Infection with HHV-6A mutants lacking telomeric repeats did not reproduce this phenotype. HHV-6A/B infections lead to increased expression of three shelterin genes, TRF1, TRF2 and TPP1. TRF2 was recruited to VRC and binding to the HHV-6A/B telomeric repeats demonstrated by chromatin immunoprecipitation and ELISA. Lastly, the HHV-6A IE2 protein colocalized with shelterin proteins at telomeres during infection. In summary, HHV-6A/B infections results in an excess of telomeric repeats that stimulates the expression of shelterin genes. TRF2 binds to viral telomeres during infection and localizes with HHV-6A IE2 protein. Our results highlight a potential role for shelterin complex proteins and IE2 during infection and possibly during integration of HHV-6A/B into host chromosomes.

## Introduction

Human herpesvirus-6A (HHV-6A) and HHV-6B are two distinct beta herpesviruses with different epidemiological and biological characteristics (1). HHV-6B is a ubiquitous virus that infects nearly 100% of world population and is the etiological agent of roseola infantum, an infantile febrile illness characterized by high fever with occasional skin rash (2). HHV-6B is also a concern in hematopoietic stem cell and solid organ transplant recipients with frequent reactivation and medical complications (3). Pathological and epidemiological data on HHV-6A remain scarce.

The viral genomes of HHV-6A/B are composed of a unique segment of approximately 143 kbp flanked at both extremities with identical and directly repeated (DR) termini of approximately 9 kbp each. Each DR contains two regions with repeated TTAGGG telomeric sequence that play a role in the ability of these viruses to integrate their genomes into human chromosomes (4) and reviewed in (5, 6). The number of telomeric repeats within each DR ranges from 15 to 180 copies in clinical isolates (7-10). Although HHV-6 integration can occur in several distinct chromosomes, it invariably takes place in the telomeric/sub-telomeric regions of chromosomes (11-14). When integration occurs in a gamete, the viral genome can be inherited resulting in individuals carrying a copy of the viral genome in every cell, a condition called inherited chromosomally integrated HHV-6 (iciHHV-6) (15). Approximately 1% of the world population is considered iciHHV-6^+^ (reviewed in (6)). The consequences of iciHHV-6 are not well defined but a recent study indicates that iciHHV-6^+^ individuals are at greater risks of developing angina pectoris (16).

The ends of mammalian chromosomes are composed the telomeres consisting in 5kb to 15 kb of telomeric repeats (TTAGGG) followed by a 200 +/- 75 nucleotide TTAGGG single-stranded 3’-overhang (17). The telomeres function as a buffer zone to avoid instability and loss of genetic information. With each cell division, the extremities of the chromosomes are incompletely replicated due to the end replication problem (18). As a consequence, the telomeres shorten after every cell division until they reach a minimal threshold length, which triggers DNA damage activation via the ATR (ataxia telangiectasia and Rad3 related) or the ATM (ataxia telangiectasia mutated) pathway ultimately leading to apoptosis or senescence (reviewed in (19, 20)). In the absence of telomere elongating processes, such as expression of telomerase or activation of the alternative lengthening of telomere (ALT) pathway, somatic cells are therefore capable of a limited number of replication cycles.

To prevent activation of DNA damage recognition pathways, chromosome ends are protected by binding the shelterin complex at telomeric repeats (19). The shelterin complex folds telomeric DNA into a secondary structure called the T-Loop, preventing the recognition of the telomere extremity as a double-strand break (DSB) (21). The shelterin complex is made of six proteins: TRF1, TRF2, TPP1, RAP1, TIN2 and POT1. TRF1 and TRF2 both form homodimers that bind directly to the double-strand TTAGGG repeats in a sequence-specific manner (22-24). TRF2 represses activation of the ATM pathway (25) and plays an essential role in end-to-end chromosome fusions mediated by the non-homologous end-joining (NHEJ) pathway (26, 27). POT1 binds to the single-strand section of the telomeres and protects the telomeres against activation of the ATR pathway (26, 28-30).

Certain viruses are reported to affect telomeres in different ways. For example, infection by herpes simplex virus type 1 (HSV-1) alters telomere integrity in several ways, including transcriptional activation of TERRA, loss of total telomeric DNA, selective degradation of TPP1, reduction of telomere-bound shelterin and accumulation of DNA damage at telomere (31). Telomere remodeling is presumed to be required for ICP8-nucleation of pre-replication compartment that stimulates HSV-1 replication (31). The Epstein-Barr virus (EBV) LMP1 protein was reported to downmodulate the expression of TRF1, TRF2 and Pot1 shelterin genes resulting in telomere dysfunction, progression of complex chromosomal rearrangements, and multinuclearity (32, 33). The impact of HHV-6A/B infection of telomere biology is currently unknown. Considering that telomeres are preferred sites for HHV-6A/B integration, it is essential to understand the dynamic processes occurring during the early phases of infection to gain insights into the integration mechanisms. In the present study, we analyzed the impact of HHV-6A/B infections on shelterin complex homeostasis and determined whether its members would associate with viral DNA during infection. We report for the first time that some shelterin proteins expressions are upregulated during HHV-6A/B infection and that TRF2 is recruited to the viral replication compartment and associates with viral DNA during infection.

## Materials and Methods

### Cell lines and viruses

U2OS cells (American Type culture collection (ATCC), Manassas, VA, USA) were cultured in Dulbecco’s modified Eagle’s medium (DMEM, Corning Cellgro, Manassas, VA, USA) supplemented with 10% Nu serum (Corning Cellgro), non-essential amino acids (Corning Cellgro), HEPES, sodium pyruvate (Multicell Wisent Inc., St-Bruno, Québec, Canada) and plasmocin 5 μg/ml (InvivoGen, San Diego, CA, USA). HeLa and MCF-7 cells (ATCC) were cultured in the same medium supplemented with 10% fetal bovine serum (FBS) (Thermo Fisher) instead of Nu serum. MOLT-3 (ATCC, CRL-1552), HSB-2 (ATCC, CCL-120.1), both human T lymphoblastic cell lines, were cultured in RPMI-1640 (Corning Cellgro) supplemented with 10% Nu serum (Corning Cellgro), HEPES and plasmocin 5 μg/ml (InvivoGen). J-JHAN cells infected with HHV-6A mutants (ΔTMR and ΔimpTMR) (4) were cultured in RPMI-1640 supplemented with 10% FBS. SUP-T1 cells were cultured in RPMI-1640 supplemented with 10% FBS. HHV-6B (Z29 strain) and HHV-6A (GS strain) were propagated on MOLT-3 and HSB-2 cells respectively, as previously described (34).

Plasmids

IE2 expression vectors (WT and Δ1290-1500) were previously described (35). pLPC-MYC-hTRF1 (Addgene plasmid # 64164) (36) and pLPC-MYC-hPOT1 (Addgene plasmid#12387) (37) were a gift from Titia de Lange and obtained through Addgene. pLKO human shTRF2 was previously described (38). The pSXneo 135(T2AG3) was a gift from Titia de Lange (Addgene plasmid # 12402) (39).

Western blots

Cells were resuspended in Laemlii buffer and boiled for 5 minutes. Samples were loaded and electrophoresed through a SDS-polyacrylamide gel. Samples were transferred onto PVDF membranes and processed for western blot using rabbit anti-TRF2 (Novus Biologicals), rabbit anti-IE1 (34), and mouse anti-tubulin antibodies (Abcam). Peroxydase-labeled goat anti-rabbit IgG and peroxydase-labeled goat anti-mouse IgG were used as secondary antibodies. The Bio-Rad Clarity ECL reagent was used for detection.

### IF-FISH and microscopy

Immunofluorescence (IF) combined with fluorescence in situ hybridization (FISH) was performed as previously described (40). U2OS cells were seeded at 5 x 10^4^ cells per well in 6-well plates over coverslips, cultured 24 hours and infected with HHV-6A or HHV-6B at a multiplicity of infection (MOI) of 5 for 4 hours. Cells were then washed with PBS and cultured in media for a set period of time. Cells were fixed with 2% paraformaldehyde. HeLa cells were treated the same way but seeded at 3.5 x 10^4^ cells per well. MOLT-3 and HSB-2 cells were infected at a MOI of 1 and cultured for a set period of time before being deposited on a 10-well microscope slide, dried and fixed in acetone at −20°C for 10 minutes. The following primary antibody were used: rabbit-α-IE1-Alexa-488 (34), mouse-α-IE2-Alexa-568 (Arsenault et al, 2003, JCV), mouse-α-P41 (NIH AIDS Reagent Program), rabbit-α-TRF2 (NB100-56694, Novus Biologicals), rabbit-α-53BP1 (H-300, Santa Cruz Biotechnology), mouse-α-γH2AX (Ser139, clone JBW301, EMD Millipore) and mouse-α-PML (PG-M3, Santa Cruz Biotechnology). Secondary antibodies used were goat-α-rabbit-Alexa-488, goat-α-rabbit-Alexa-594, goat-α-mouse-Alexa-488 and goat-α-mouse-Alexa-594 (Life Technologies). FISH was performed using a PNA probe specific to the telomeric sequence (CCCTAA)_3_ (TelC-Cy5, PNA BIO).

Slides were observed using a spinning disc confocal microscope (Leica DMI6000B) and analyzed using the Volocity software 5.4.

To compare TRF2 expression in uninfected and HHV-6-infected cells, cells were dually stained with HHV-6 IE2 protein and TRF2. The relative TRF2 fluorescence in IE2- and IE2+ individual cell was then determined using the ImageJ software. TRF2 expression levels were compared using unpaired student t-test with Welch’s correction.

Telomere restriction fragment (TRF) analysis.

DNA from uninfected and HHV-6A/B infected cells was isolated using QIAamp DNA blood isolation kits as per the manufacturer’s recommendations. Five µg of DNA were digested overnight with RsaI and HinfI followed by electrophoresis through agarose gel and southern blot hybridization. The telomeric DNA probe was obtained following digestion of the pSXneo 135(T2AG3) vector with EcoRI and NotI, gel purification of the 820 bp fragment and ^32^P-labeling by nick translation. After hybridization and washes, the membrane was exposed to X-ray films.

### RNA isolation and RT-qPCR

HSB-2 and MOLT-3 cells (10^7^ cells) were incubated in a 50 mL tube and infected with HHV-6A or HHV-6B at a MOI of 0,25. After 4h at 37°C, cells were washed and placed in a 25 cm^2^ culture flask at 500 000 cells per ml in complete media. A portion of the cells were harvested on day 1, 2, 3, 5 and 7 post-infection. Total RNA was isolated and processed by reverse transcriptase quantitative PCR (RT-QPCR) as previously described (41). The cDNAs obtained were analyzed by TaqMan qPCR using Rotor-Gene Q apparatus (Qiagen) and Rotor-Gene Multiplex PCR Kit reagent (Qiagen) using validated HHV-6-IE1- and GAPDH-specific primers and probes (41). cDNAs were analyzed for TRF1, TIN2, TPP1 using specific primers and SyBr green as described (42). TRF2, POT1, RAP1 were analyzed using the following primers:

TRF2 FWD: 5’-GTACCCAAAGGCAAGTGGAA-3’ TRF2 REV: 5’-TGACCCACTCGCTTTCTTCT-3’

POT1 FWD: 5’-TGAAGTTCTTTAAGCCCCCA-3’ POT1 REV: 5’-AGCCTGTGAAAGCGAACAAT-3’ RAP1 FWD: 5’-GCCACCCGGGAGTTTGA-3’ RAP1 REV: 5’-GGGTGGATCATCATCACACATAGT-3’

The C_t_ value of genes from infected cells was compared to the value of uninfected cells and normalized with the GAPDH cellular gene.

### Flow cytometry

HSB-2 cells were mock-treated or infected with HHV-6A (U1102) at a MOI of 0.5. After 48 hours, 1 x10^6^ cells/assay were fixed, permeabilized and processed for detection of HHV-6 antigens (p41 (9A5D12 from NIH AIDS Reagent Program) and gp102 (7A2 from NIH AIDS Reagent Program)) and TRF2 (Novus Biologicals) using the intracellular fixation and permeabilization buffer kit (eBiosciences, San Diego, CA, USA). One μg of antibody was used per assay. Cells were analyzed by flow cytometry using a FACS scan apparatus and CellQuest software.

### ChIP and dot blot

The experiments were made using the Pierce Magnetic ChIP Kit (Thermo Scientific) according to the manufacturer’s instructions with a few modifications. Equal quantities of HSB-2 and MOLT-3 cells were used for all samples (4 x 10^6^ cells/sample). Cross-linking lasted 10 minutes at RT. Two μl of diluted MNase (1:10) were added to each sample for MNase digestion. Before sonication, an aliquot was saved for normalization purpose (input). Sonication was made with a Branson Sonifier 450, with an Output Control set at 1. Each sample was sonicated with five pulses of 20 seconds, each pulse followed by a 20 seconds incubation on ice. Before immunoprecipitation, samples were incubated with magnetic beads alone for one hour at 4°C before discarding the beads. The immunoprecipitation was performed using 2 μL of normal rabbit IgG (negative control) and 4 μg of rabbit anti-TRF2 antibody (NB100-56694, Novus Biologicals) with an overnight incubation at 4°C. Protein A agarose beads were added for 1h at 4°C followed by three washes. The DNA was eluted in 50 μL of DNA column elution solution.

Eluted DNA was analyzed by dot blot hybridization using telomeric probe ((CCCTTA)_4_ probe) or HHV-6 probe (DR6) while the input was analyzed using an Alu probe. The DNA was first denatured for 10 minutes at room temperature in 0.25 N NaOH and 0.5 M NaCl. Samples were then serially diluted in 0.1 X SSC and 0.125 N NaOH, on ice, loaded onto nylon membrane, neutralized in 0.5 M NaCl and 0.5 M Tris-HCl pH 7.5 and crosslinked using UV irradiation. Membranes were pre-incubated in Perfecthyb Plus hybridization buffer (Sigma-Aldrich) for 2h at 60°C before addition of 1 x 10^6^ CPM/ml of ^32^P-labeled probes. Hybridization was carried out for 16h at 60°C. Membrane was washed twice with 2X SSC-1% SDS, twice with 1X SSC-1% SDS and once with 0.5X SSC-1% SDS at 60°C, for 15 minutes each. Membrane was then exposed to X-ray films at −80°C.

### Cloning and purification of MBP-TRF2

The pLPC-NMYC TRF2 was a gift from Titia de Lange (Addgene plasmid # 16066). The TRF2 coding sequence was excised from pLPC-NMYC TRF2 vector using with BamHI and XhoI and cloned in frame with the MBP coding sequence of the pMAL-C2 vector (New England Biolabs) using BamHI and SalI enzymes. MBP and MBP-TRF2 proteins were expressed in BL21 DE3 RIL bacteria and purified by affinity chromatography, as described (43).

### Electrophoretic mobility shift assay (EMSA)

EMSA was performed essentially as described (43). In brief, recombinant proteins (MBP and MBP-TRF2) were incubated with double-stranded (ds) non-telomeric or telomeric labeled probes in 20 µl of the following reaction buffer: 20 mM Hepes-KOH pH 7.9, 150 mM KCl, 1 mM MgCl_2_, 0.1 mM EDTA, 0.5 mM DTT, 5% glycerol and 0.1 mg/ml BSA. For competition experiments, 10-1000 fold excess unlabeled ds non-telo or telomeric probes were included in the reaction buffer. After a 30 minute incubation at room temperature, 2 µl of loading dye were added and the samples were electrophoresed through a non-denaturing 5% acrylamide:bis (29:1) gel. After migration, the gels were dried and exposed to X-ray films at −80°C.

### Detection of TRF2 binding to HHV-6 telomeric sequence

The wells of a 96-well ELISA plate were coated with 25 ng MBP or 50 ng of MBP-TRF2 proteins by overnight incubation at 4°C in pH 9.0 carbonate buffer. After rinsing, 1% BSA was added to block non-specific sites. Twenty-five nanograms of HaeIII-digested digoxigenin-labeled HHV-6A DNA (HaeIII cuts the viral genome 289 times) in EMSA reaction buffer were added. For competition experiments, 2.5 or 5.0 pmoles of non-telomeric or telomeric dsDNA were added 15 minutes prior to the addition of HHV-6A DNA. The plate was incubated for 2h at room temperature (RT). After 3 washes with TBS-0.1% Tween-20 (TBS-T), peroxidase-labeled mouse anti-DIG antibodies were added to each well for 1h at RT. After 3 additional TBS-T washes, TMB substrate was added and the reaction allowed to develop for 15 minutes before addition of 50µl of 2N sulfuric acid. Absorbance was measured at 450 nm.

## Results

### Telomeric sequence accumulation during HHV-6A infection

Telomeres help protect against the loss of genetic information due the linear DNA end replication problems encountered during each cell division. Telomeric repeats are the binding sites of six proteins referred to as the shelterin complex that prevent induction of a DDR at chromosome ends. Interestingly, the extremities of the HHV-6A/B genomes also contain stretches of telomeric (TTAGGG)_n_ repeats that vary in number between 15 and 180 (7-10) (Figure 1A). Using fluorescent *in situ* hybridization (FISH), we first studied the accumulation of telomeric sequences during active HHV-6A infection. Hybridization of mock-infected HSB-2 cells with a telomeric probe resulted in the detection of many discrete punctate telomeric signals corresponding to chromosome telomeres (Figure 1B). In contrast, a mixture of small and enlarged telomeric signals were observed during HHV-6A infection. At late stages of infection when viral genome replication is abundant, very intense telomeric signals were detected. These telomeric signals likely correspond to replicating virus genomes as they localize with the IE2 protein that is presumed to associate with viral DNA (44) (Figure 1B, bottom panels). Similar results were observed in HHV-6B-infected cells (Figure 1C). Using ddPCR we have quantified the number of viral DNA copies/cell at 96h post-infection and assuming all cells are productively-infected, results indicate that between 15,000 and 18,000 viral DNA molecules/per cell are present during active infection. Considering that 116 and 80 TTAGGG repeats/genome are respectively present in HHV-6A and HHV-6B (7), between 750 000 and 2 million telomeric repeats are present in HHV-6A/B-infected cells compared to 125 000 telomeric repeats in uninfected cells (assuming 8kbp/telomere) (Figure 1D). Such increase in telomeric signals was also monitored by terminal restriction fragment (TRF) analysis. As shown in figure 1E, uninfected cells displayed telomeres lengths ≥2kbp. In addition to the cellular telomeric signal observed, HHV-6A/B infected cells displayed abundant signals that were smaller in size (<2kbp) and with much stronger in intensity, representing viral telomeric signals.

**Figure 1:**
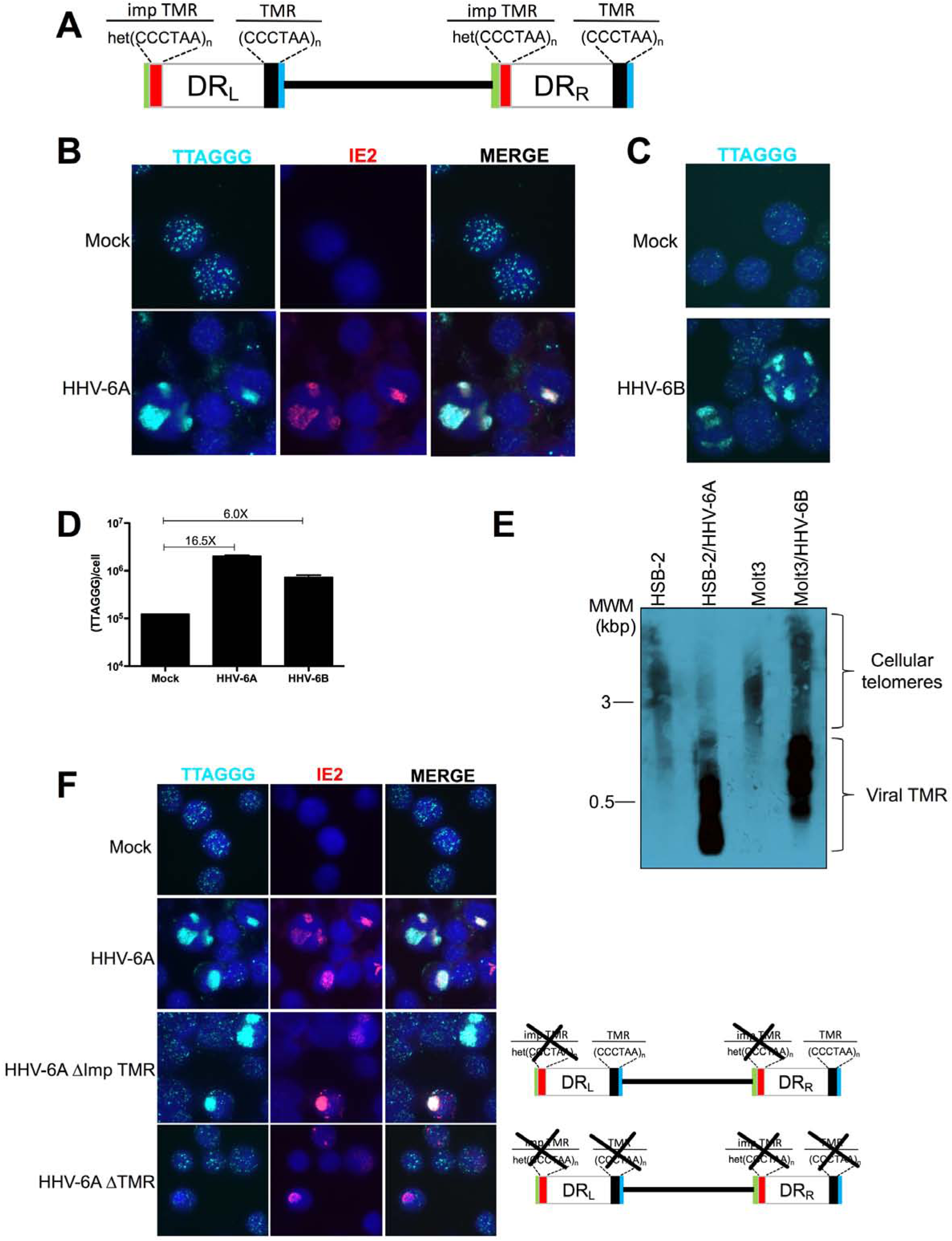
Accumulation of viral telomeric signals during HHV-6A infection. A) Schematic representation of the HHV-6A/B genome. The unique (U) region of the HHV-6A/B genome (140 kbp) is flanked by two direct repeat sequences (10-13 kbp) referred to as DR_L_ and DR_R_. The DRs contain perfect (CCCTAA)_n_ and imperfect (het(CCCTAA)_n_ telomeric sequences. The genome is not drawn to scale. B) HSB-2 cells were infected with HHV-6A. After 5 days of infection, cells were processed for IF-FISH to detect HHV-6A IE2 protein (red) and telomeres (cyan) using a telomeric probe. Nuclei were stained with DAPI. C) J-JHAN cells were infected with a recombinant HHV-6A or mutants lacking the imperfect telomeric repeats (ΔimpTMR) or lacking all telomeric repeats (ΔTMR). After several days of infection, cells were processed for IF-FISH to detect HHV-6A IE2 protein and telomeres (cyan). Nuclei were stained with DAPI.

The accumulation of telomeric sequences was confirmed using J-JHAN cells infected with a recombinant HHV-6A. As with HSB-2 cells, mock-infected J-JHAN showed typical telomeric staining (Figure 1F, first row). J-JHAN cells productively infected with HHV6-A demonstrated large telomeric signals that colocalized with the viral IE2 protein (second row). To demonstrate that the increase telomeric signals observed originates from viral DNA, we made use of HHV-6A mutants lacking either only the imperfect telomeric repeats (ΔimpTMR) or all telomeric repeats (ΔTMR) (4). Infection with the ΔimpTMR mutant still resulted in a strong and patchy telomeric signals (Figure 1F, third row). In contrast, telomeric hybridization signals in ΔTMR-infected J-JHAN cells were similar to those observed in uninfected J-JHAN cells (Figure 1F last row), confirming that telomeric sequences within the viral genome were responsible for the increased telomeric signals observed.

### Modulation of the shelterin complex expression during HHV-6A/B infection

Telomeres are bound by a series of 6 proteins referred to as the shelterin complex. Considering the increase in telomeric sequences detected during active HHV-6A/B infections, we surmised that shelterin expression is likely to be modulated during infection. Total RNA was isolated at varying time points post HHV-6A/B infections and RT-qPCR was performed using primers specific for each of the shelterin genes as well as the telomerase gene. The cellular *GAPDH* gene was used for normalization. As showed in figures 2A-F, HHV-6A infection of HSB-2 cells led to increased expression of TRF1, TRF2, RAP1 and TPP1 mRNAs with statistical significance (p<0.05) observed on day 7 post-infection. Variations in POT1, and TIN2 mRNA levels were only minimal. Similarly, HHV-6B infection of MOLT-3 cells caused a significant increase in TRF2 and TPP1 mRNAs levels relative to uninfected control cells (Figures 2H-M). Other shelterin genes were not significantly modulated. HHV-6 infection was monitored by assessing U90 gene expression (Figure 2G).

**Figure 2:**
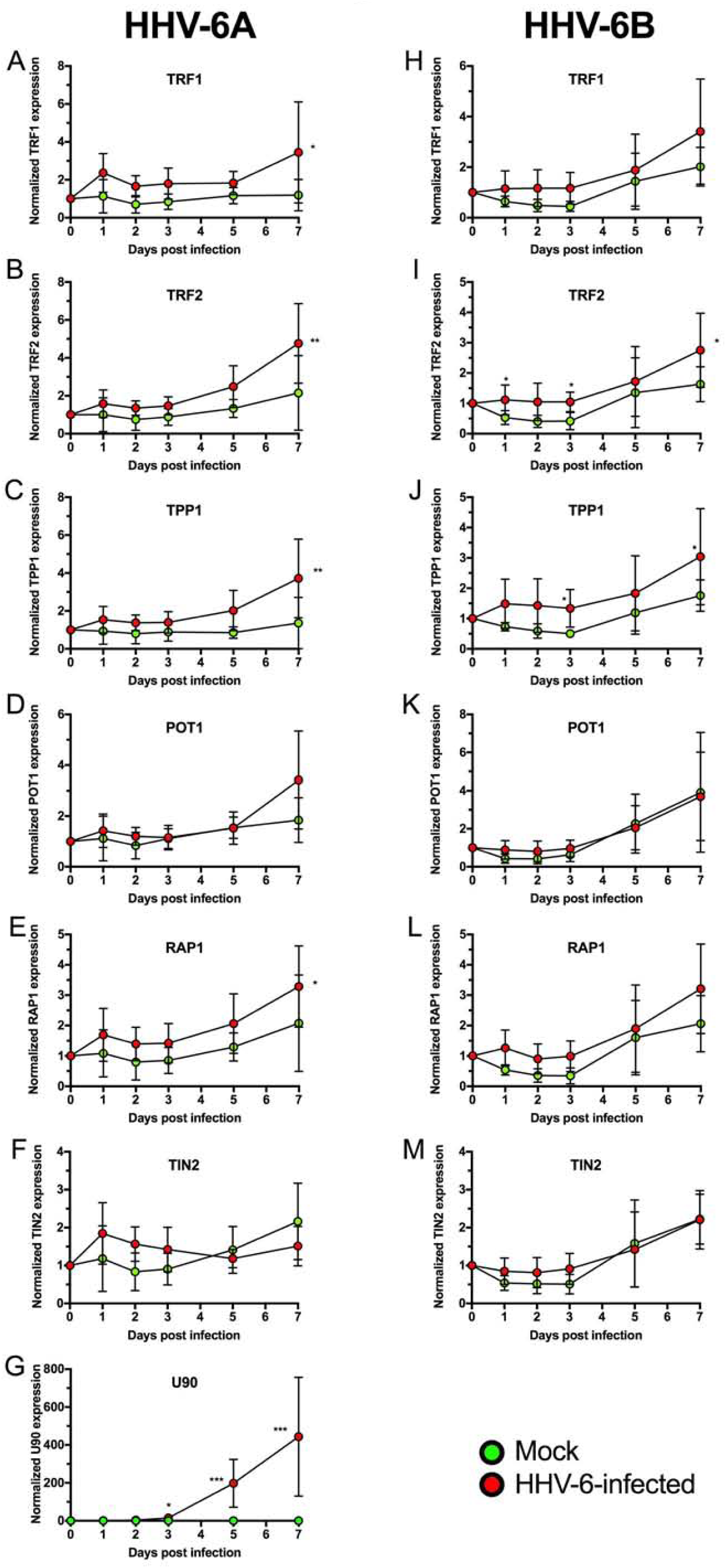
Kinetics of shelterin genes expression during HHV-6A/B infection. HSB-2 cells (A-G) and Molt3 cells (H-M) were respectively infected with HHV-6A or HHV-6B. At various time post infection, total RNA was extracted and analyzed by reverse transcriptase QPCR for *TRF1, TRF2, POT1, RAP1, TIN2, TPP1, GAPDH and U90* genes expression. Shelterin genes expression was normalized relative to *GAPDH* gene expression while U90 was analyzed to demonstrate infection. Results represent data from 4-6 independent experiments expressed as mean +/-SD gene expression relative to that of uninfected cells. *p<0.05.

We next investigated whether these changes in shelterin mRNA levels would translate in increased protein expression. We infected HSB-2 cells with HHV-6A and analysed intranuclear TRF2 expression by flow cytometry. HHV-6A-infected cells were identified by detection of P41 or gp102 viral proteins expression. The mean TRF2 fluorescence intensity (MFI) in uninfected cells varied between 261 and 210 (Figures 3A-B). In HHV-6A-infected cells, two cell populations were observed. Infected ones, expressing P41 or gp102, and uninfected bystander cells. As shown, HHV-6A-infected cells expressed TRF2 at higher levels (MFI of 385 and 376) relative to uninfected cells within the same population (MFI of 204 and 259). Similar results were obtained in HHV-6B-infected Molt3 cells with a MFI TRF2 of 314 in infected cells relative to MFI TRF2 of 180 and 117 in uninfected or p41^-^ cells, respectively (Figure 3C).

**Figure 3:**
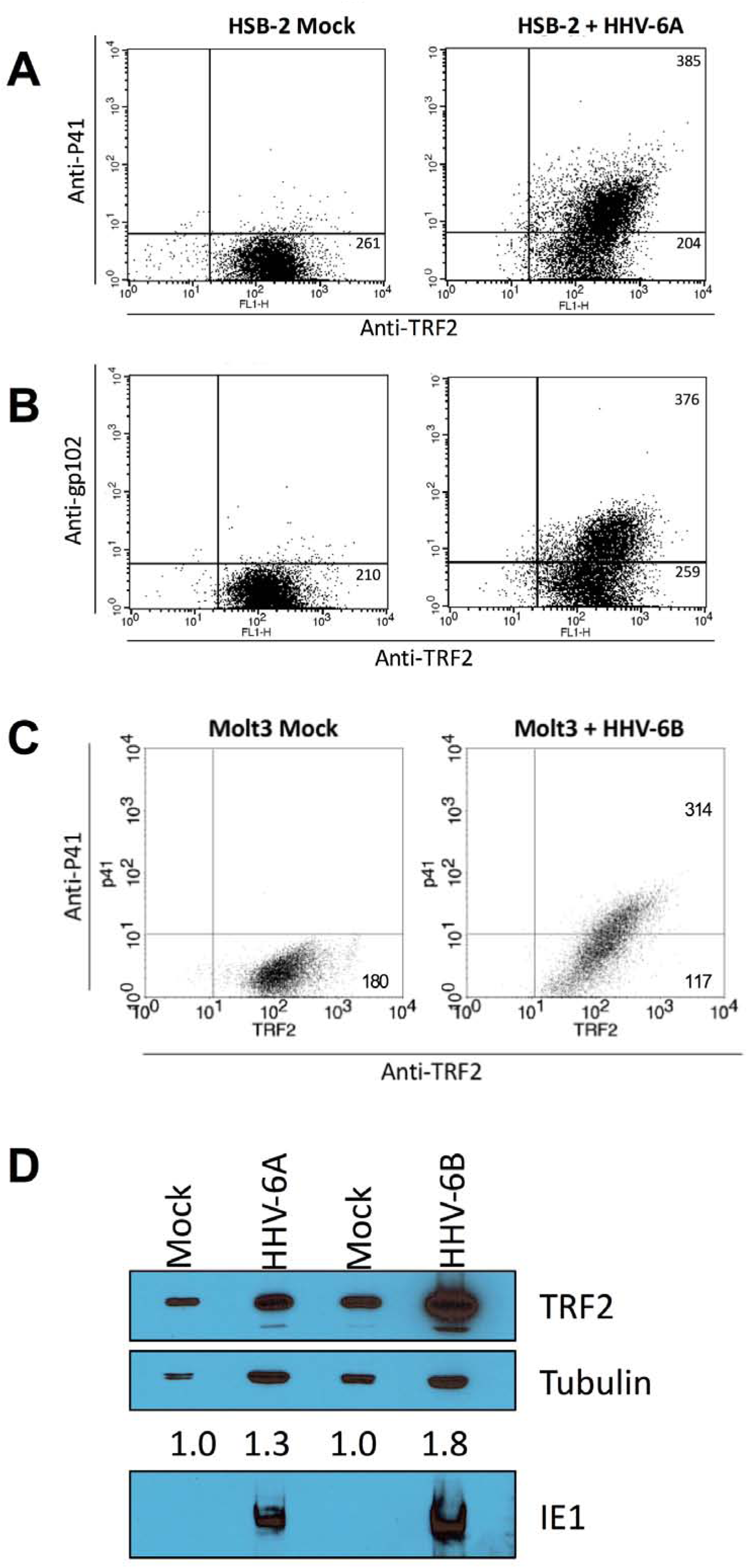
TRF2 expression during productive HHV-6A/B infections. Mock, HHV-6A- or HHV-6B-infected cells were analyzed for TRF2 expression by flow cytometry. Uninfected and 5 days old HHV-6A-infected HSB-2 cells (A-B) and HHV-6B-infected Molt3 cells (C) were fixed, permeabilized and stained for TRF2, P41 and gp102 proteins expression. Numbers in the top and bottom left quadrants indicate mean relative TRF2 fluorescence intensities. Results are representative of two independent experiments. D) Western blot analysis of TRF2 expression in HHV-6A/B infected. Tubulin was used as loading controls and IE1 to demonstrate HHV-6A/B infection. Numbers represent TRF2 expression levels relative to mock-infected cells after normalization with tubulin.

TRF2 expression levels in mock-infected and HHV-6A/B infected cells was also assessed by western blot analysis. As shown in figure 3D, compared to mock-infected cells, HHV-6A and HHV-6B-infected cells had TRF2 protein expression levels that were increased 1.3X and 1.8X, respectively. These results (Figures 3A-D) confirm the mRNA data and indicate that TRF2 is expressed at higher levels in HHV-6A/B-infected cells.

T cell lines such as HSB-2 and Molt3 are highly susceptible to HHV-6 replication and typically used for HHV-6 propagation. As most cells are lytically infected and subsequently killed, HHV-6A/B integration in such cells does not occur frequently. We therefore determined whether TRF2 expression would also be modulated in semi-permissive cells, such as U2OS that we and others routinely use to study HHV-6A/B integration (45). TRF2 expression in HHV-6A-infected U2OS cells was measured using confocal microscopy. After 24h, 48h and 72h post-infection, individual cells were analyzed for TRF2 expression. Infected cells were distinguished from uninfected bystander cells using the anti-IE2 antibody (Figure 4A). Fluorescence was quantified using ImageJ software (Figure 4B). The results obtained indicate that starting at 24h post-HHV-6A infection, TRF2 is expressed at significantly higher levels in infected cells than bystanders or uninfected cells (p ≤ 0.02). No significant difference in TRF2 expression was detected between bystander and mock-treated cells. In summary, these results indicate that expression of TRF2 increases during HHV-6A/B-infections.

**Figure 4.**
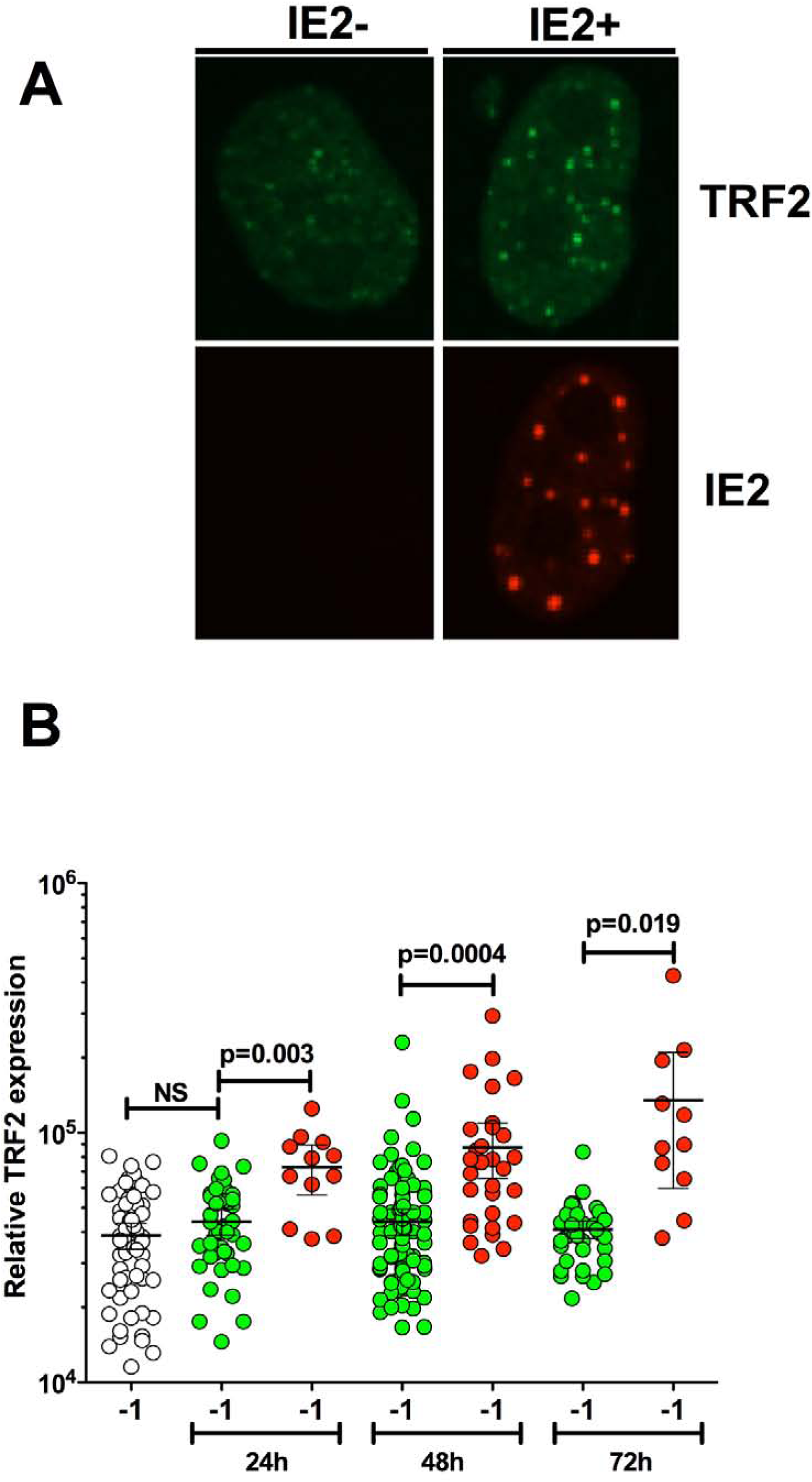
Increased TRF2 expression in HHV-6A-infected U2OS cells. U2OS cells were infected with HHV-6A and analyzed for TRF2 and IE2 expression at 24h, 48h and 72h post-infection by dual color immunofluorescence. A) Representative TRF2 and IE2 expression in bystander and IE2 expressing cells at 48h post infection. B) Mean relative TRF2 expression ± SD in uninfected (white), IE2-(green-uninfected bystander) or IE2+ (red-infected) cells at 24h, 48h and 72h post infection. Each symbol represents the relative TRF2 expression from a single cell.

### Binding of TRF2 to viral telomeric sequences

Telomeres are protected by the shelterin complex of which TRF1 and TRF2 bind directly to TTAGGG repeats; however, it remains unknown if shelterin proteins bind to viral telomeric sequences in the context of the HHV-6A/B genomes. To study TRF2 binding to viral TMRs, a recombinant MBP-TRF2 protein was generated. To validate that MBP-TRF2 was functional and capable of binding telomeric DNA, we performed EMSA. MBP-TRF2 efficiently bound dsDNA with telomeric sequences causing a mobility shift (figure 5A). MBP alone did not bind the telomeric probe. The specificity of MBP-TRF2 binding was confirmed by a competition with excess unlabeled telomeric and non-telomeric oligonucleotides. Excess (100-1000 fold) of unlabeled telomeric oligonucleotides efficiently competed with labeled telomeric probes. No such competition was observed with excess non-telomeric oligonucleotides. Lastly, no binding of MBP or MBP-TRF2 was observed using non-telomeric labeled probes (Figure 5B).

**Figure 5:**
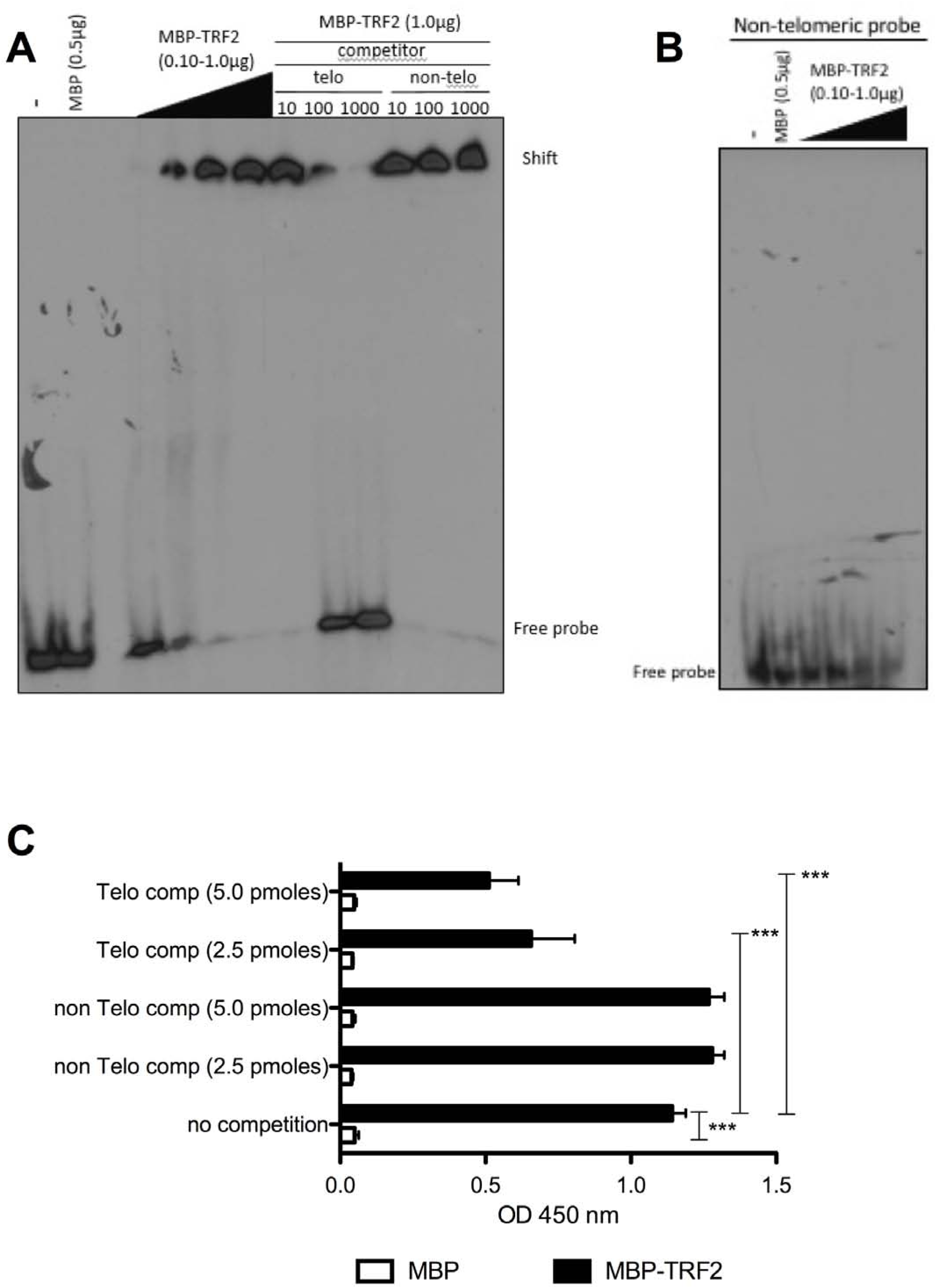
Binding of TRF2 to HHV-6 viral DNA. Recombinant MBP or MBP-TRF2 were incubated with ^32^P-labeled telomeric dsDNA (A) and binding was assessed by EMSA. Excess of unlabeled telomeric and non-telomeric dsDNA were added as competitors. Samples were migrated on non-denaturing acrylamide gel, dried and exposed to X-ray films. B) Recombinant MBP or MBP-TRF2 were incubated with ^32^P-labeled non-telomeric dsDNA and binding was assessed by EMSA. C) Recombinant MBP and MBP-TRF2 were coated to the wells of a 96 well-plate and incubated with *HaeIII* digested DIG-labeled HHV-6A DNA (25 ng/condition) in the presence or absence of competitors. After washing, bound DNA was quantified by adding peroxidase-labeled anti-DIG antibodies and substrate. Results are expressed as mead absorbance +SD of triplicate values. Experiment is representative of two additional experiments. *** P<0.001.

After validation of the specific binding to telomere sequences of the recombinant MBP-TRF2 protein, we next determined if MBP-TRF2 is able to bind to HHV-6 TMR DNA. To study this, DIG-labeled HHV-6A-BAC DNA was digested with the HaeIII enzyme that cuts on both sides of the viral TMR and more than 250 times in the viral genome. MBP and MBP-TRF2 coated plates were incubated with the mixtures of DNA fragments (25 ng) and DNA binding was measured using anti-DIG antibodies. MBP did not bind viral DNA, in contrast to MBP-TRF2 that efficiently bound viral DNA (Figure 5C). Specificity of MBP-TRF2 binding to viral TMR was confirmed through successful competition with unlabeled ds oligonucleotides containing telomeric motifs (Telo comp) but not by ds oligonucleotides with non telomeric motifs (non Telo comp). Our *in vitro* binding assay revealed that the recombinant MBP-TRF2 efficiently binds to viral DNA at TMR.

To validate these results, colocalization of TRF2 with viral DNA during infection was studied next. As reported previously (45), HHV-6A/B infection of U2OS is abortive in most cells with little or no viral DNA replication observed. In such cells, the HHV-6A IE2 protein was dispersed throughout the nucleus in several independent foci (Figure 4). However, in a minority of cells, IE2 displays a patchy appearance reminiscent of viral replication compartments (VRC) (Figure 6A). The presence of IE2 to VRC was confirmed by co-staining cells for the HHV-6 DNA processivity factor P41 that is known to associate with the viral DNA polymerase (46, 47). P41 and IE2 associated with large diffuse telomeric signals that represent viral TMRs (Figure 6A). Colocalization of TRF2 at VRC was studied next. As shown in figure 6B, in HHV-6A-infected U2OS cells with VRC, TRF2 colocalized with IE2 along with diffuse telomeric signals. In infected cells where VRC were not detected (most cells), the presence of diffuse telomeric signals and “patchy” IE2 was not observed (Figure 6B, last row). Despite the absence of VRC, TRF2 and IE2 often colocalized during infection. To determine if ectopically-expressed IE2 would colocalize to telomeres and TRF2 in the absence of other viral proteins or viral DNA, U2OS were transfected with an empty vector or an IE2 expression vector and cells were analyzed by IF-FISH. IE2 displayed a punctate nuclear distribution, with while most of IE2 colocalized with endogenous TRF2 (Figure 6C). Whether IE2 requires its C-terminal DNA-binding domain (DBD) for telomeric localization was studied next. Cells expressing an IE2 mutant lacking the DBD mutant was found to localize at telomeres as efficiently as WT IE2 (figure 6D).

**Figure 6.**
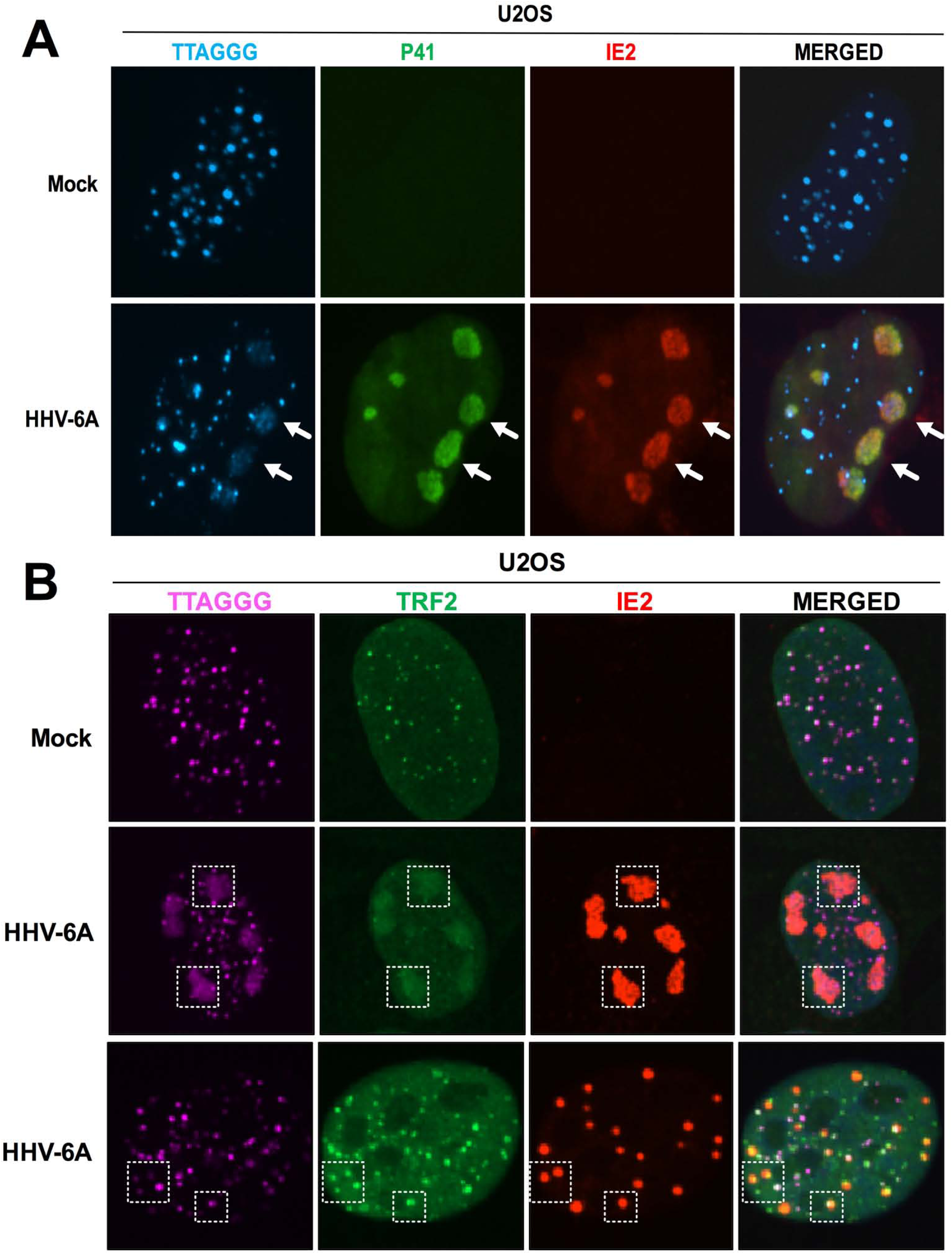
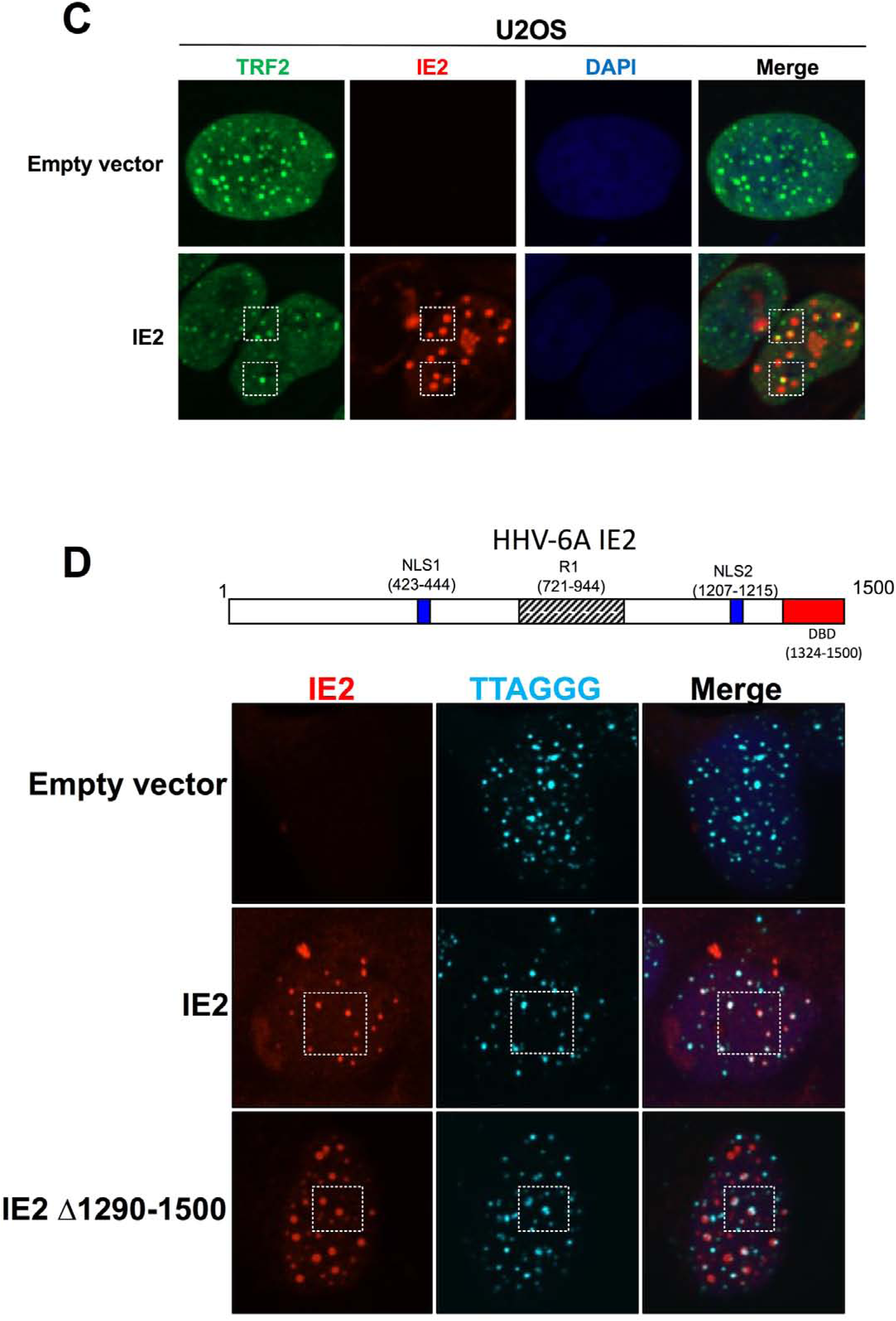
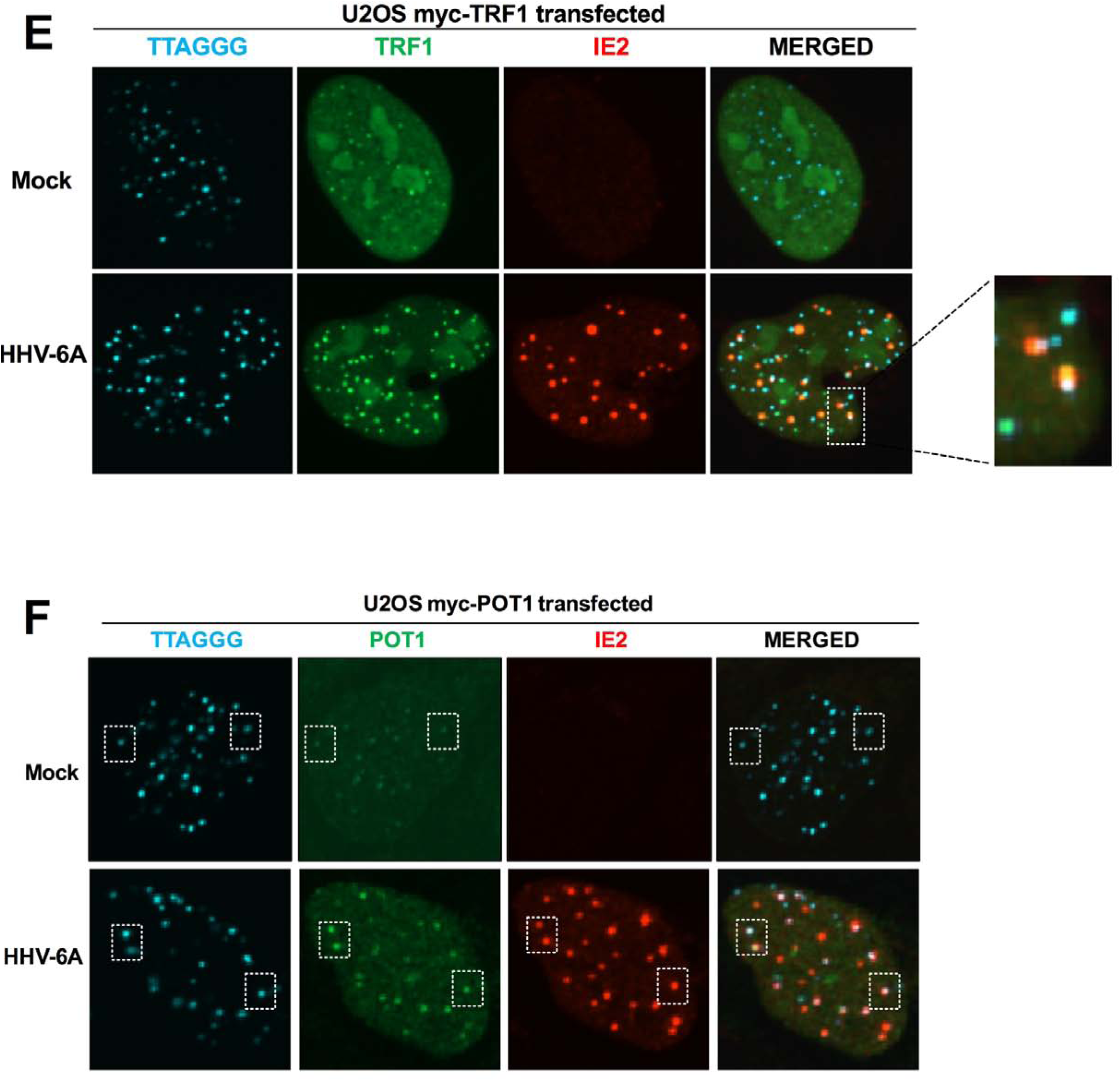
Colocalization of shelterin complex proteins and HHV-6A IE2 protein at viral and cellular telomeres. A) U2OS cells were infected for 48h with HHV-6A after which cells were processed for IF-FISH. Telomeres were labeled in blue, p41 in green and IE2 in red. These images demonstrate colocalization of IE2 with P41, a viral protein that associates with viral DNA during infection, and diffuse telomeric signals (arrows). B) Telomeres were labeled in magenta, TRF2 in green and IE2 in red. The panels in the middle row show images of cells productively infected (minority of cells) with HHV-6A. Large diffuse telomeric signals (viral replication compartments) where TRF2 and IE2 accumulates (rectangles) are represented. The panels in the third row represent infected cells that do not actively replicate viral DNA with TRF2 and IE2 colocalizing (dashed squares) at distinct telomeres. C) Colocalization of HHV-6A IE2 protein at telomeres in the absence of viral DNA. U2OS cells were transfected with an empty vector or an IE2 expression vector. Forty-eight hours later cells were processed for dual color immunofluorescence. TRF2 was labeled in green and IE2 in red. Examples of IE2 colocalizing with TRF2 are presented (dashed squares). D) U2OS cells were transfected with WT IE2 or IE2 Δ1290-1500 expression vectors. Forty-eight hours later cells were processed for IF-FISH. Telomeres were labeled in cyan, IE2 in red and nuclei in blue. Examples of IE2 colocalizing with TRF2 are presented (dashed squares). E) Uninfected and HHV-6A-infected U2OS cells were transfected with an empty vector or a myc tagged TRF1 expression vector. Forty-eight hours later cells were processed for IF-FISH. Telomeres were labels in cyan, TRF1 in green and IE2 in red. Examples of TRF1 localizing at telomeres (dashed squares) in uninfected cells are shown in the top row. Examples of IE2 colocalizing with TRF1 and telomeres in infected cells are presented in the bottom row (dashed squares). F) Uninfected and HHV-6A-infected U2OS cells were transfected with an empty vector or a myc tagged POT1 expression vector. Forty-eight hours later cells were processed for IF-FISH. Telomeres were labels in cyan, POT1 in green and IE2 in red. Examples of POT1 localizing at telomeres (dashed squares) in uninfected cells are shown in the top row. Examples of IE2 colocalizing with POT1 and telomeres in infected cells are presented in the bottom row (dashed squares).

We next investigated if HHV-6A IE2 protein would colocalize with other shelterin proteins during infection. U2OS were transfected with myc-tagged TRF1 and POT1 expression vectors, infected with HHV-6A and analyzed by IF-FISH. Both TRF1 and POT1 colocalized with telomeres in control cells as expected (Figures 6E-F). In HHV-6A-infected cells, IE2 was found to partially colocalize with both TRF1 and POT1 at telomeres.

So far, the results obtained indicate that IE2 colocalizes with shelterin complex at telomeres. In addition, TRF2 colocalizes with VRC during infection, suggesting that telomeric sequences within HHV-6 DNA are likely recognized and bound by TRF2. To provide additional support to this hypothesis, we performed TRF2 ChIP in HHV-6A and HHV-6B productively-infected cells. Uninfected cells were used as negative controls. Using equal amounts of starting material, DNA-bound by TRF2 was immunoprecipitated (IP) using anti-TRF2 antibodies and the DNA analyzed by dot blot hybridization. TRF2 efficiently bound telomeric sequences in both uninfected and HHV-6A- and HHV-6B-infected cells (Figures 7B-E). In addition, a stronger telomeric signal was observed in infected cells relative to uninfected cells. To discriminate between telomeres of cellular and viral origin, the TRF2 immunoprecipitated DNA was hybridized with the DR6 probe, corresponding to regions adjacent (1.5kbp) to the TMR in the virus genome (refer to Figure 7A). As shown, the DR6 probe preferentially bound to DNA isolated from HHV-6-infected cells (Figures 7B-E). As negative control, DNA was immunoprecipitated with an irrelevant mouse anti-IgG. As positive control, DNA was IP using anti-RNA PolII antibodies and analyzed by qPCR for GAPDH promoter DNA (not shown). These results indicated that during infection, HHV-6A and HHV-6B TMRs are physically bound by TRF2.

**Figure 7.**
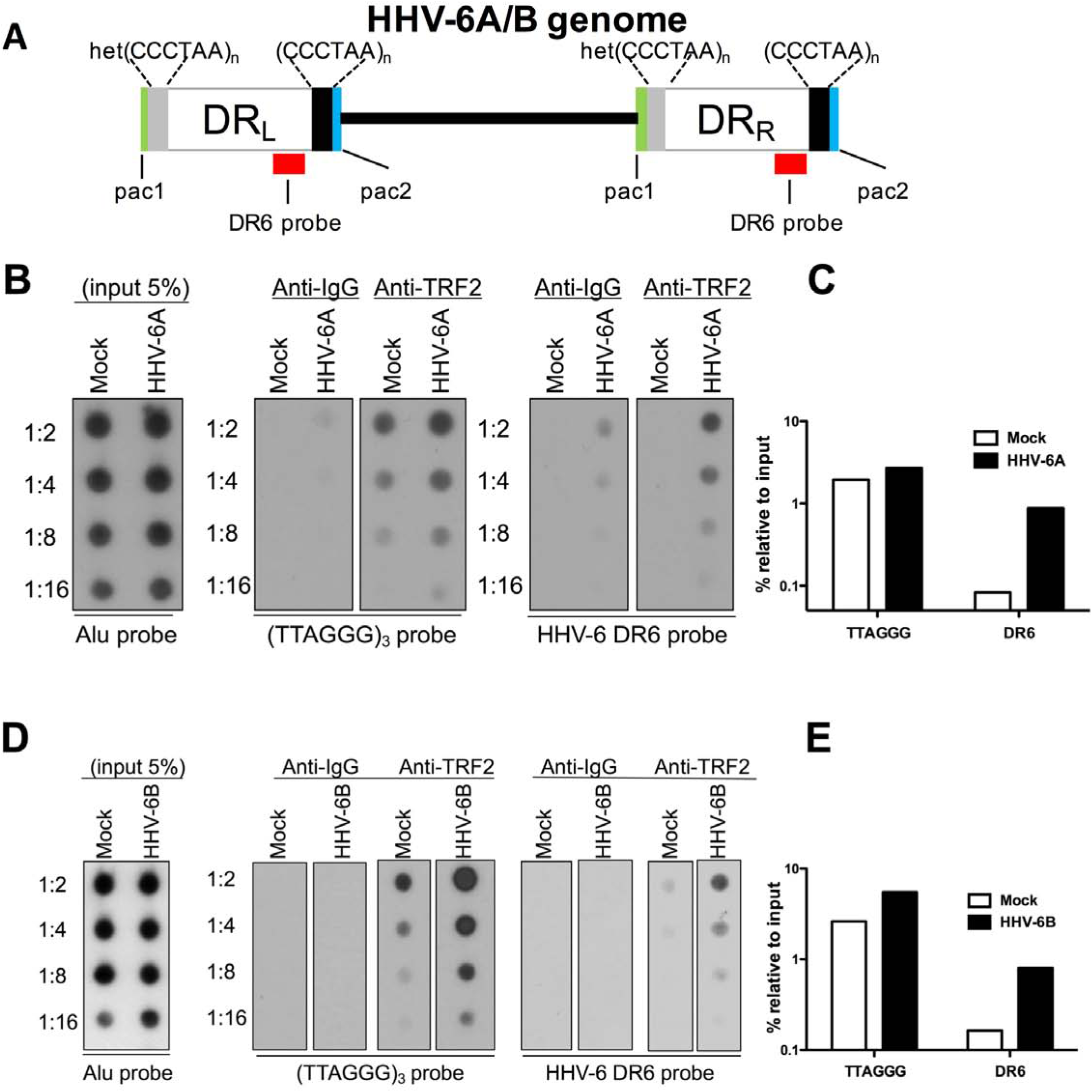
Binding of TRF2 to viral DNA during HHV-6A/B infection. A) Schematic representation of the HHV-6A/B genome. The DR6 probe used for hybridization is shown in red. Uninfected and HHV-6A-infected HSB-2 cells (B-C) or uninfected and HHV-6B-infectd Molt3 cells (D-E) were analyzed for TRF2 binding to viral DNA using ChIP. The input was hybridized with Alu probe to assess quantity of starting material. Anti-IgG (negative control) or TRF2 antibodies were used for immunoprecipitation. Eluted DNA was serially diluted and hybridized with ^32^P-labeled telomeric (TTAGGG)_3_ or HHV-6 (DR6) probes. After hybridization the membranes were washed and exposed to X-ray films. The quantity of TRF2 bound to telomeric and viral DNA is measured relative to the input. Results are of 3 independent experiments.

In summary, these assays provide evidence that TRF2 binds the telomeric motifs present in HHV-6A/B DNA.

### TRF2 is not required for IE2 localization at viral replication compartment

Considering that HHV-6A IE2 colocalizes with TRF2 at cellular telomeres (Figures 6B-C) and with TRF2 at VRC (Figure 6B), we hypothesized that TRF2 might influence IE2 localization. We generated an U2OS cell line carrying a doxycycline (Dox)-inducible shRNA targeting TRF2 mRNA. Incubation of cells with Dox for 7 days resulted in TRF2 knockdown (Figure 8A). In the absence of Dox, TRF2 and IE2 were found to localize at VRC following HHV-6A infection (Figure 8B, top row). Upon TRF2 knockdown, IE2 still localized efficiently to VRC, indicating that TRF2 is dispensable for IE2 localization at VRC. To confirm that TRF2 knockdown was sufficient to induce a phenotype, the DDR at telomeres was assessed. Cells expressing TRF2 (-DOX) expressed little or no phospho 53BP1, a DDR marker (Figure 8C). In contrast, mock-infected and HHV-6A-infected cells in which TRF2 knockdown was induced (+DOX) showed robust 53BP1 expression at telomeres, suggesting that TRF2 expression was below the levels required to maintain telomere protection (Figure 8C).

**Figure 8.**
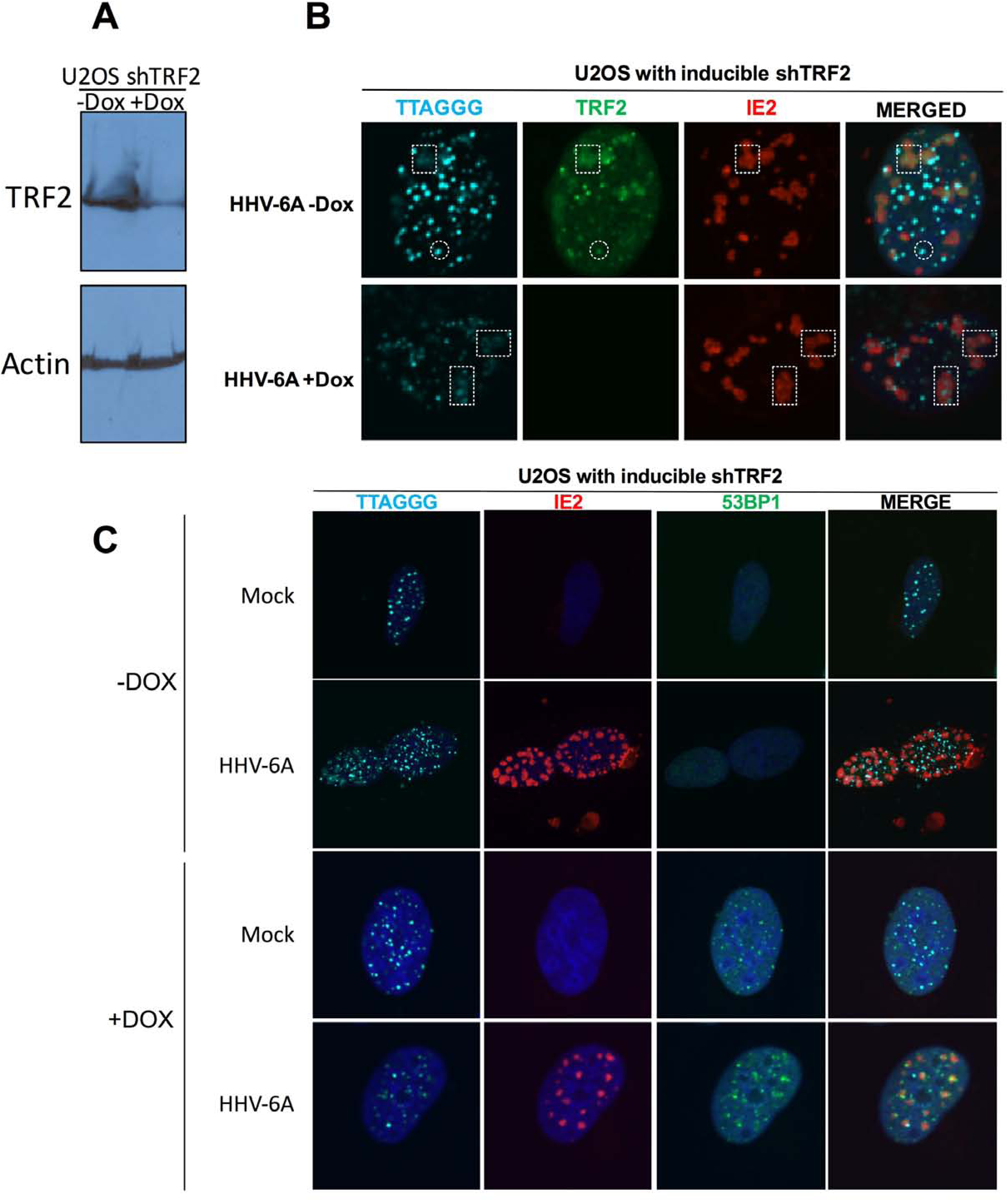
IE2 localized to VRC in the absence of TRF2. U2OS cells were transduced with a lentiviral vector coding for a Dox inducible shRNA against TRF2. Transduced cells were selected with puromycin for a week. A) Half of the cultures was treated with Dox for seven days to induce TRF2 knockdown (KD), as determined by western blot. B) Control (-Dox) and TRF2 KD (+Dox) cells were infected with HHV-6A for 48h and processed for IF-FISH. Telomeres were labeled in cyan, TRF2 in green and IE2 in red. As show in the–Dox condition, TRF2 colocalized with IE2 as well as diffuse (dashed square) and punctate (dashed circle) telomeric signals. In the +Dox condition, TRF2 KD was confirmed with IE2 colocalizing with diffuse telomere signals (dashed squares). C) DDR at telomeres as a consequence of TRF2 knockdown. U2OS cells were treated or not with Dox and infected with HHV-6A as in figure 6B. Cells were then processed for IF-FISH. Telomeres were labeled in cyan, IE2 in red, 53BP1 (as marker of DDR) in green and nuclei in blue.

### Importance of TRF2 during productive viral infection

Considering that TRF2 colocalize and interacts with viral DNA during HHV-6A/B infections, we determined if TRF2 knockdown would affect HHV-6A/B DNA replications. SUP-T1 cells, susceptible to both HHV-6A and HHV-6B infections, were transduced with a Dox-inducible shTRF2 encoding lentiviral vector. After selection, cells were treated or not with Dox for 20 days after which TRF2 expression levels were monitored by western blots. TRF2 expression was significantly reduced after 20 days of Dox (Figure 9A). Control (-Dox) and TRF2 KD SUP-T1 cells (+Dox) were infected with HHV-6A or HHV-6B. Infections were allowed to proceed, and intracellular DNA collected at varying time points and analyzed by ddPCR to assess HHV-6A/B DNA copies. The relative quantity of viral DNA was very similar between cells having normal or reduced TRF2 levels (Figures 9B-C), suggesting that TRF2 depletion had only marginal effects on HHV-6A/B DNA replication.

**Figure 9.**
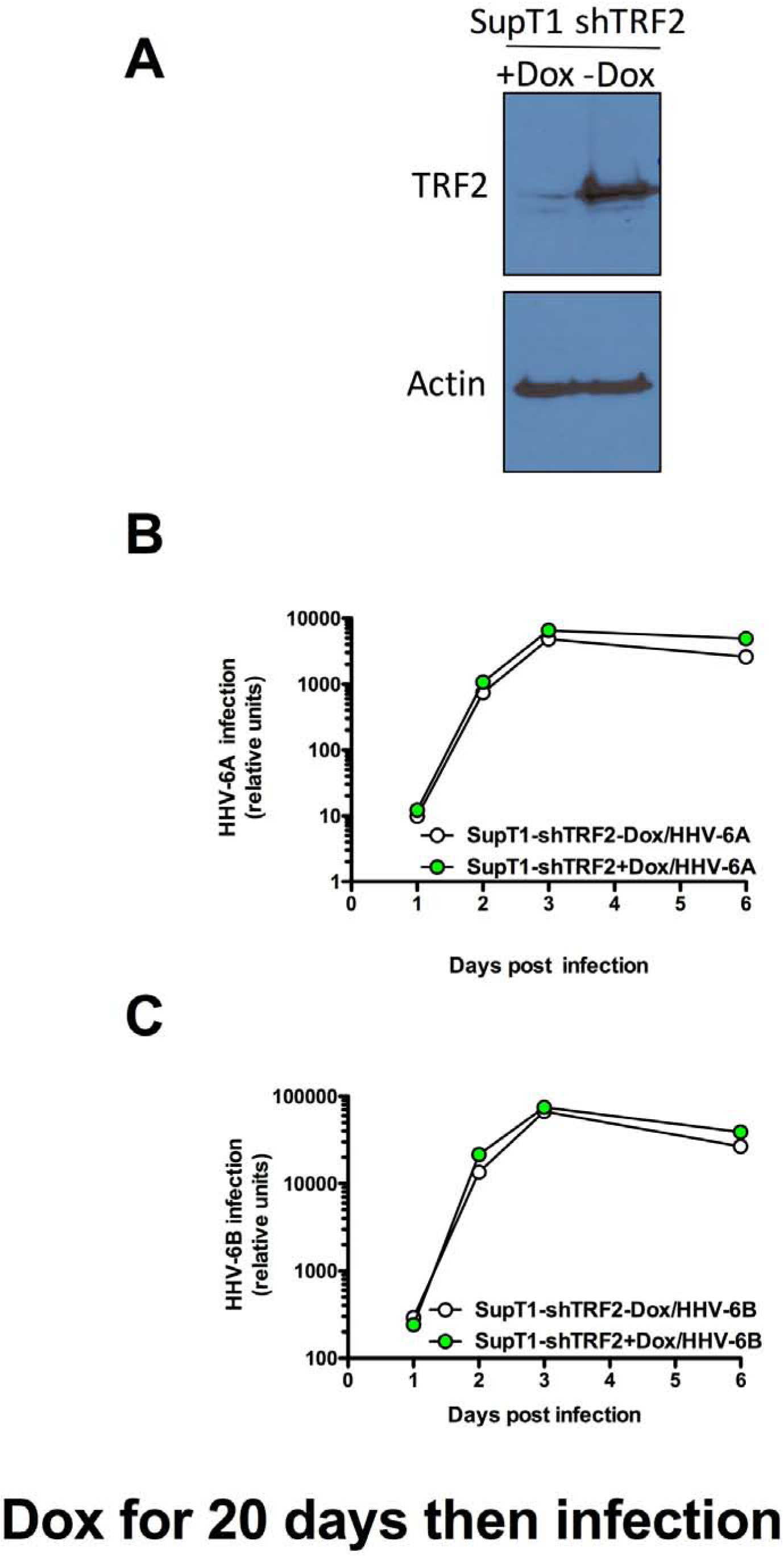
Knockdown of TRF2 does not affect HHV-6A/B replication. SUP-T1 cells were transduced with a lentiviral vector coding for a Dox inducible shRNA against TRF2. A) Transduced cells were selected with puromycin for two weeks. TRF2 knockdown (KD) was induced by adding Dox to the culture medium for three weeks and confirmed by western blot. B-C) Control (-Dox) and TRF2 KD (+Dox) SUP-T1 cells were infected with HHV-6A (B) or HHV-6B (C). Whole cell DNA was isolated at various time points and the relative number of HHV-6A/B genomes determined and normalized against cellular DNA.

## Discussion

Telomeres serve to protect chromosomes from the loss of genetic information. Each chromosome contains several hundreds, even thousands, of tandemly repeated TTAGGG hexamers. Each time a cell divides approximately 150 nucleotides are lost due the end replication problem (48). When telomeres get short, these are extended either by the telomerase enzyme complex (49) or alternative lengthening mechanisms (50, 51). On the other hand, when telomeres get excessively long, proteins such as TZAP, can trim the excess telomeres (52). Mechanisms sensing the length of telomeres are therefore present in cells to control telomere length. In the present study, we report that during HHV-6A/B infection, the number of TTAGGG repeats increases significantly (Figures 1B-C). The increase in telomeric sequences originates from the viral genomes that contain between 15 and 180 TTAGGG repeats at each viral extremity (7-10). A HHV-6A mutant lacking these telomeric sequences does not reproduce this phenotype. Using ddPCR we calculated the number of viral DNA molecules to be in excess of 10,000 per cell resulting in an increase in telomeric repeats, in a productively-infected cell, that is 6 to 16 times higher than that of an uninfected cell. The cells responded to this increase in neo telomeric sequences by turning on TRF1, TRF2, TPP1 and RAP1 genes expression. No increase in POT1 or TIN2 was observed. The lack of POT1 induction does not come as a surprise considering that POT1 binds to single-stranded telomeric motifs and no such ssDNA is generated by the viral DNA during infection. At the protein level, TRF2 overexpression in HHV-6A/B infected cells was demonstrated by IFA and FACS. The fact that TRF2 upregulation was observed only in HHV-6A/B infected cells indicates a direct consequence of infection rather than potential paracrine effects. TRF2 was found to localize at viral replication compartments along with the HHV-6A IE2 and P41 proteins, the latter being a viral DNA polymerase processivity factor (47). The IE2 protein is a large nuclear protein (circa 1500 amino acids) that behaves as a promiscuous transactivator in gene reporter assays (35, 44). We have previously reported that truncation of the C-terminus abolishes IE2’s transactivating potential (35). Recently, the crystal structure of the IE2 C-terminus revealed that it contains dimerization, DNA-binding and transcription factor binding domains explaining the importance of this region for IE2’s functions (53). Although IE2 localizes at VRC during infection, whether it binds viral DNA *per se* remains to be demonstrated. Considering that IE2 localizes at HHV-6A ΔTMR VRC, suggests that IE2 does not preferentially bind telomeric DNA repeats. Furthermore, considering that in the absence of viral DNA IE2 localizes at cellular telomeres suggest a potential affinity for certain shelterin complex proteins. Of interest, the IE2 C-terminus core structure resembles those of the gammaherpesvirus factors EBNA1 of Epstein-Barr virus (EBV) and LANA of Kaposi sarcoma-associated herpesvirus (KSHV) (53), involved in binding to viral DNA (54, 55). Deletion of the IE2 DNA binding domain had no impact on IE2 localization at telomeres (Figure 6D), further strengthening the hypothesis the IE2 interacts with telomere binding proteins.

Using ChIP, we could demonstrate that TRF2 associates with viral DNA during infection. Furthermore, using recombinant TRF2 and BAC viral DNA, we could show that TRF2 binds to viral DNA in the absence of other factors. Binding could be efficiently competed with dsDNA containing telomeric repeats indicating TRF2 binding to viral telomeric sequences. By being attracted to HHV-6A/B TMRs, TRF2 may leave telomeres unprotected, leading to instability or damage repair. The increase in TRF2 production by the infected cell may compensate for the potential reduction of TRF2 at cellular telomeres. Alternatively, infected cells may respond to the presence of numerous viral TMRs present by turning on *TRF2* gene expression, which is supported by our data. Furthermore, the lack of noticeable DDR at telomeres (data not shown) during infection argues in favor of the latter explanation.

Shelterin protein binding to DNA of other viruses has been reported previously. Binding of TRF2, TRF1 and Rap1 to EBV *oriP*, that contains three TTAGGGTTA motifs, was reported to modulate EBV DNA replication. TRF2 also interacts with EBNA1, an EBV protein essential for episomal maintenance and replication (56). While TRF2 and Rap1 promote the replication at *oriP*, TRF1 inhibits it (56-58). TRF2, together with KSHV LANA protein bind to the latent origin of replication. Such region does not contain the TTAGGG motif and binding to this region of the viral DNA likely involves a yet to be identified protein (59). Unlike EBV, the expression of a dominant negative TRF2 does not affect KSHV DNA replication. In that regard, our results indicating that TRF2 silencing had no impact on HHV-6A/B DNA replication are similar to those of Hu et al (59).

During infection, many viruses provoke a DNA damage response, either because their unprotected genome is recognized as damaged DNA or because of viral proteins triggering a damage signal. While several viruses have ways to evade the DDR pathways, some have developed strategies to make use of the cellular DNA repair proteins to their advantage. Cellular DNA repair proteins have been observed in viral replication compartment in various cases and can be helpful or even necessary for completion of the infection (60). During Epstein-Barr virus (EBV) infection, the proteins involved in the ATM pathway checkpoint and HR repair are found in replication compartments (61). The use of the DDR machinery by EBV likely increases the possibility of molecular events, stimulating the damage signals causing instability and promoting carcinogenic transformations. Whether viruses can use the DDR proteins in chromosomal integration is controversial, but some studies have suggested it (60). One example is the Adeno-associated virus (AAV) that uses the cellular NHEJ mechanism for its site-specific integration (62). HHV-6A/B chromosomal integration is not fully understood but it appears probable that these viruses integrate by HR between the virus’ TMRs and the cellular telomeres. The integration occurs solely in telomeres and it has been shown that the telomeric sequences within the HHV-6A genome are essentials for efficient integration into chromosomes (4). Furthermore, the integrated virus has been sequenced and its orientation and missing sections are compatible with an integration by HR between the viral TMRs and the telomeres (11, 63, 64). Whether TRF2 plays a role in HHV-6A/B integrations remains to be shown. Considering the importance of TRF2 in maintaining telomere integrity, it is very challenging to demonstrate its role in integration as our current *in vitro* assays typically require culturing of the cells for a month (45). Depletion of TRF2 for extended times results in chromosome fusions and subsequent cell death, preventing us from conducting such experiments.

In summary, we showed that during HHV-6A/B infection, the number of telomeric repeats increases significantly. Such an excess of unprotected telomeric repeats stimulates the expression of shelterin genes. The shelterin protein TRF2 binds to viral telomeres during infection and localize with HHV-6A IE2 protein at viral replication compartments. Our results highlight a potential role for shelterin complex proteins and IE2 during infection and possibly during integration of HHV-6A/B into host chromosomes.

## ACKNOWLEDGMENTS.

We acknowledge the Bioimaging platform of the Infectious Disease Research Centre, funded by an equipment and infrastructure grant from the Canadian Foundation Innovation (CFI). This work was funded by a Canadian Institutes of Health Research grants (MOP_123214 and PJT_156118) awarded to LF. SGG and VC are recipients of fellowships from the Fonds de Recherche Québec-Santé.

## AUTHOR CONTRIBUTIONS

Conceived experiments: S.G.G. and L.F. Performed experiments: S.G.G., A.G., VC, L.F. Contribution of key reagents and methodology: B.B.K.,D.J.W., E.L.D. Data analysis: A.G., S.G.G., and L.F., Writing: S.G.G. and L.F. Manuscript revision: S.G.G., A.G., B.B.K., E.L.D., D.J.W., V.C., L.F.

